# Genomic MET amplification occurs early in NF1-related malignant peripheral nerve sheath tumor (MPNST) progression and is a potent therapeutic target

**DOI:** 10.1101/199000

**Authors:** Jacqueline D. Peacock, Matthew G. Pridgeon, Elizabeth A. Tovar, Curt J. Essenburg, Megan Bowman, Zachary Madaj, Julie Koeman, Jamie Grit, Rebecca D. Dodd, Diana M. Cardona, Mark Chen, David G. Kirsch, Flavio Maina, Rosanna Dono, Mary E. Winn, Carrie R. Graveel, Matthew R. Steensma

## Abstract

Malignant Peripheral Nerve Sheath Tumors (MPNSTs) are highly resistant sarcomas that occur in up to 13% of individuals with Neurofibromatosis Type 1 (NF1). Genomic analysis of longitudinally collected tumor samples in a case of MPNST disease progression revealed early hemizygous microdeletions in *NF1* and *TP53*, with concomitant amplifications of *MET*, *HGF*, and *EGFR*. To examine the role of MET in MPNST progression, we developed mice with enhanced MET expression and NF1 ablation (*NF1*^*fl/KO*^;*lox-stop-loxMET*^*tg/+*^;*Plp-creERT*^*tg/+*^; referred to as NF1-MET). NF1-MET mice express a robust MPNST phenotype in the absence of additional mutations. A comparison of NF1-MET MPSNTs with MPNSTs derived from NF1^KO/+^;p53^R172H^;Plp-creERT^tg/+^ (NF1-P53) and NF1^KO/+^;Plp-creERT^tg/+^ (NF1) mice revealed unique Met, Ras, and PI3K signaling patterns. To investigate the therapeutic potential of MET inhibition among tumorgrafts derived from the respective MPNST models, we tested the highly selective MET inhibitor, capmatinib. NF1-MET MPNSTs were uniformly sensitive to MET inhibition whereas only a small subset of NF1-P53 and NF1 MPNSTs were inhibited. These results confirm that MET activation is sufficient for Schwann cell dedifferentiation into MPNSTs in the context of NF1 deficiency. RAS-MET signal interactions may be an important driver of MPSNT disease progression.

## Introduction

Neurofibromatosis type 1 (NF1) is caused by germline mutations in the NF1 gene and is the most common single-gene disorder affecting ∽1 in 3,000 live births (1,2). Approximately 8-13% of individuals with NF1 will develop malignant tumors, most commonly Malignant Peripheral Nerve Sheath Tumors (MPNSTs) (3). NF1-related MPNSTs are highly aggressive sarcomas that frequently metastasize and have five-year survival rates ranging from 20-50% (4-8). The mainstay of treatment is surgical resection when possible with consideration of chemotherapy and radiation therapy in select cases. Even though chemotherapy may initially stabilize disease, early responses are typically followed by a rapid evolution of chemoresistance and metastasis (9).

The NF1 gene encodes neurofibromin, a GTPase-activating protein that regulates RAS (including HRAS, NRAS, and KRAS) and loss of NF1 leads to deregulated RAS signaling. The RAS signaling node activates multiple kinase effector cascades, including the RAF – MEK – ERK pathway. NF1-related MPNSTs have been shown to arise from NF1-null myelinating Schwann cells where neurofibromin deficiency results in RAS deregulation (10-15). Plexiform neurofibromas are the benign precursor of NF1-related MPNSTs and are formed by a recruited admixture of *NF1* haploinsufficient cells (fibroblasts, mast cells, and perineurial cells) following an initial NF1-loss-of-heterozygosity event in a peripheral nerve Schwann cell (16-18). A variety of genomic alterations have been observed in the malignant transformation of benign plexiform neurofibromas into MPNSTs including deleterious mutations that impact cell cycle regulation *(CDKN2A*), apoptosis (*TP53*), tumor suppression (*PTEN, RASSF1A*) and chromatin modification (*SUZ12/PRC2* gene family) (19-21). Genes encoding receptor tyrosine kinase (RTKs) and their putative ligands are also amplified in MPNSTs, including *MET*, *HGF*, *EGFR*, *ERBB2*, *PDGFRα*, and *c-Kit*. NF1-related MPNSTs exhibit significant intertumoral heterogeneity, and it is difficult to determine the relative impact of common MPNST genomic alterations on disease progression and therapeutic response (22-24).

The oncogene MET encodes a receptor tyrosine kinase that is involved in the progression and metastasis of most solid human cancers (25-27). The pleiotropic effects of MET activation are mediated through a variety of effector pathways including dominant regulators of cellular proliferation and survival (i.e. RAS / ERK, PI3K / AKT / mTOR) and cellular motility (STAT3, Rho kinases. Several studies have implicated oncogenic MET signal activation in NF1-related MPNST disease progression. An increase in MET expression has been observed in the malignant transformation of plexiform neurofibromas into MPNSTs (28). Furthermore, high-resolution array-CGH recently identified *MET* and *HGF* gene amplifications in a significant proportion (∽30%) of NF1-related MPNSTs (24). MET phosphorylation (Tyr1234 / 35) was demonstrated in at least half of MPNSTs, and was recently proposed as a biomarker for MET-activated MPNSTs (29). In this same study, inhibition of MET / VEGFR2 / RET (cabozantinib) mitigated tumor growth in an MPNST xenograft model. Currently, it is unclear 1) whether MET activation is sufficient for malignant transformation of NF1-deficient Schwann cells to into MPNSTs, and 2) how the RAS and MET signaling pathways interact in MPNSTs.

Here, we used a longitudinal genomic analysis to identify key genetic events underlying transformation of a plexiform neurofibroma to MPNST. Our results indicate that there is positive selection for *MET* and *HGF* copy number gain early in MPNST progression that precedes accumulation of other oncogene amplifications and additional losses of tumor suppression gene function. In order to interrogate the role of MET signaling in MPNST progression and therapy response, we developed and characterized a unique mouse model of MET activation in *p53* wild type, NF1-null myelinating cells. Our hypothesis is that MET activation is sufficient to drive malignant tumorigenesis when combined with putative NF1 loss of function, and that MET-activated MPNSTs will respond to targeted MET inhibition. Our findings build upon prior work examining *NF1* loss of heterozygosity (LOH) in myelinating cells (30,31) and present a complementary model of MPNST to established models combining *NF1* loss with *p53* or *Ink4a / Arf* (19,32). Our results demonstrate that highly selective MET inhibition is effective against MPNSTs bearing a “MET addicted” signature whereby MET inhibition also mitigates downstream RAS-ERK and PI3K-AKT activation. Conversely, we show reduced effectiveness of MET inhibitor monotherapy in *Met*-amplified NF1-P53 and *Hgf -* amplified NF1 MPNSTs. These findings are attributable in part to signaling adaptations that occur early during therapy. These results expand our current understanding of the role of MET signaling in MPNST disease progression and identify a potential therapeutic niche for NF1-related MPNSTs. Moreover, these findings highlight the influence of co-occurring genomic alterations on RAS effector signaling and therapy response to tyrosine kinase inhibitors.

## Results

### Longitudinal genomic analysis of MPNST progression confirms progressive genomic instability

To investigate the genomic alterations that occur during MPNST progression, we followed the disease course of an NF1-affected adolescent male who developed a high grade, metastatic MPNST arising from a chest wall plexiform neurofibroma. Exome sequencing was performed on biopsies procured throughout disease progression: benign plexiform neurofibroma → MPNST pretreatment → MPNST posttreatment → MPNST metastasis (Figure 1A). The patient initially presented at 15 years of age with a rapidly enlarging mass in his left axilla that measured 11 x 9 x 17cm by MRI (Figure 1B, left panel). Biopsies were performed at diagnosis which verified the presence of a high grade MPNST arising within a plexiform neurofibroma (AJCC stage 3, T2bG3N0M0). The patient received neoadjuvant chemotherapy consisting of 3 cycles of ifosfamide and doxorubicin. To enhance the resectability of the tumor following neoadjuvant chemotherapy, the patient received 5000 cGy of preoperative radiation concurrently with two further cycles of ifosfamide. The combined radiation and chemotherapy reduced the size of the tumor enough to permit wide resection with chest wall reconstruction four months after diagnosis Figure 1B, middle panel). Following resection, the patient completed two additional cycles of chemotherapy with ifosfamide and doxorubicin. Surveillance imaging obtained four months after the end of therapy revealed recurrent disease in both lungs (Figure 1B, right panel) and CT-guided biopsy confirmed the diagnosis of metastatic MPNST. The recurrence was not amenable to resection and the patient succumbed to disease two years from the date of original diagnosis

**Figure 1:**
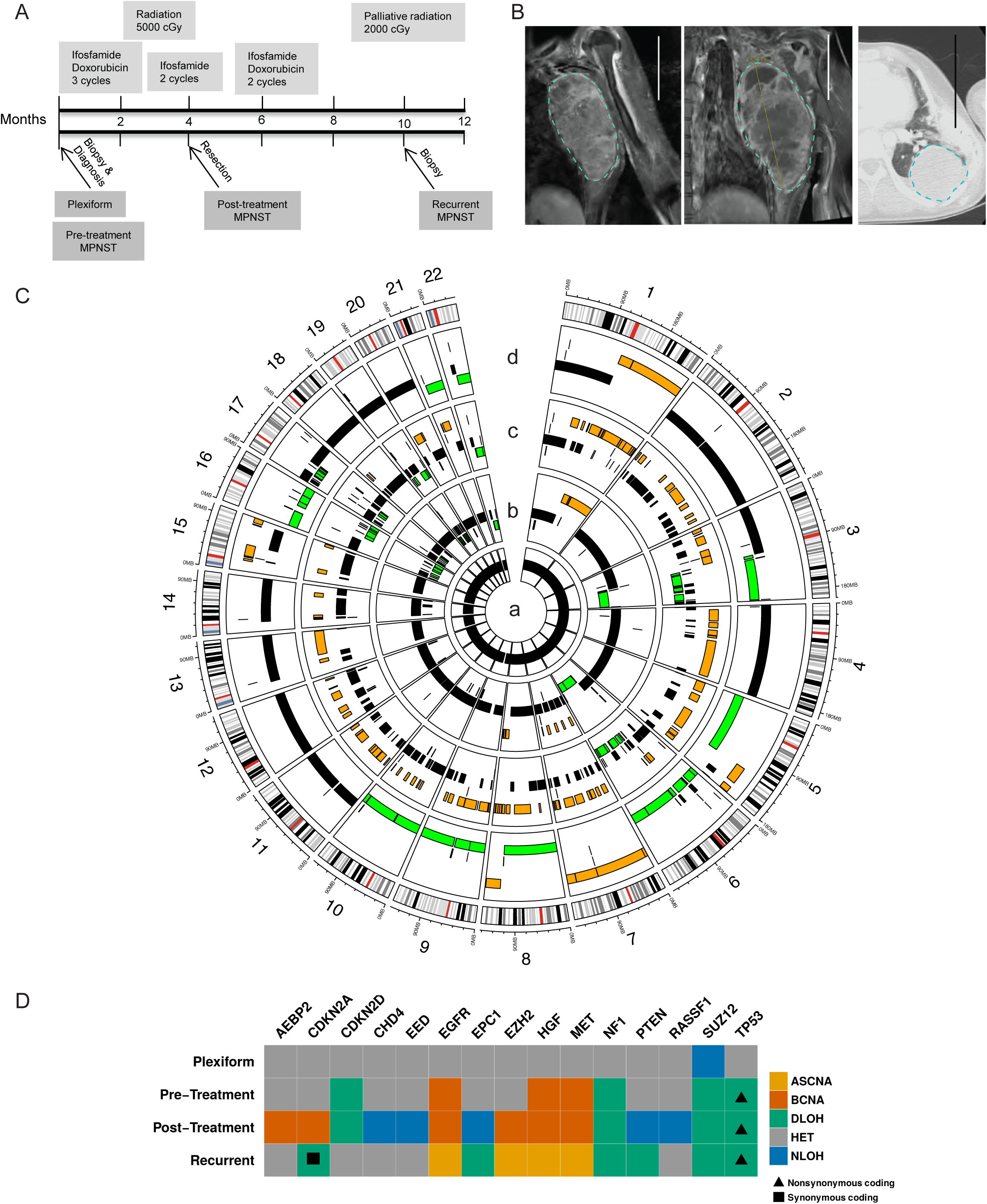
Comprehensive, longitudinal genomic analysis demonstrates targetable genetic changes early in disease progression. (A) Timeline with treatments (upper) and biopsy samples (arrows, lower; n=1 sample per time point) from an MPNST and associated plexiform neurofibroma in an adolescent male with Neurofibromatosis type 1. (B) The MPNST (indicated by blue dotted line) progression was imaged by magnetic resonance imaging at diagnosis (left panel) and post-treatment (middle panel). The recurrence was imaged by computerized axial tomography (CAT) (right panel). (C) Summary of genomic copy number changes across the four tumor samples. The plexiform neurofibroma (inner ring, a) was genomically normal. Increased copy number gains (orange), and deletions (green) throughout the progression of the pre-treatment MPNST (ring b), post-treatment MPNST (ring c) and recurrent MPNST (ring d). (D) Detailed assessment of copy number changes in known MPNST-related genes with overlaid sequence alteration data identified allele-specific copy number amplifications (ASCNA), balanced copy number amplifications (BCNA), deletion LOH (DLOH), heterozygous alleles (HET), and copy-neutral LOH (NLOH). Triangles indicate a non-synonymous coding change in TP53 (NC_000017.11:g.7673223G>C).

To evaluate the genomic alterations that occurred during MPNST progression in this case, tumor DNA was analyzed using a shotgun whole exome sequencing approach. Copy number alterations were inferred using TITAN with the patient's peripheral blood DNA serving as a normal control. The genomic alterations that were identified in each stage of MPNST progression are summarized in a Circos plot (Figure 1C). As expected, the plexiform neurofibroma DNA had few copy number variations (ring a, Figure 1C), but the pre-treatment MPNST demonstrated chromosomal amplifications (orange regions) on chromosomes 1, 7, and 8 (ring b, Figure 1C). These amplifications were maintained throughout the remaining disease progression. Additional structural alterations and site-specific amplifications accumulated throughout the course of treatment (ring c, Figure 1C). Regional amplifications on chromosome (chr) 5, chr15, a potential whole chromosomal amplification of chr7, and deletion of chr16 (ring a, Figure 1C) developed in the metastatic lesion, specifically. Several deletions (green) were also identified in the MPNST, including a region of chr17q that contains *TP53* and *NF1*. Notably, the deleted regions of chr17q as well as regions on chr3, chr6, and chr16 progressively expanded throughout disease progression. Focused analysis of known MPNST-related loci *AEBP2, CDKN2A, CDKN2D, CHD4, EED, EGFR, EPC1, EZH2, HGF, MET, NF1, PTEN,RASSF1, SUZ12,* and *TP53* was performed (Figure 1D). The SUZ12 locus was found to be altered in the plexiform neurofibroma, within a genomic segment that while diploid, is representative of a near loss of heterozygosity event on chromosome 17. Somatic mutations in SUZ12 were not observed, but a single germline intron variant was identified. Therefore, the near loss of heterozygosity is representative of a potential genomic structural change within the SUZ12 region of chromosome 17. The observed structural alterations appear to be nonrandom and favor gain of oncogenic receptor tyrosine kinases and loss of *TP53*.

The earliest observed genomic alterations in the MPNST were balanced copy number gains in *MET*, *HGF*, *EGFR*, and to a lesser degree amplification of *CDKN2D*. Changes in these specific loci preceded other sites of additional oncogene amplification or tumor suppressor loss. The number of amplified *MET*, *HGF*, and *EGFR* loci continued to increase during MPNST progression (Figure 2A). Specifically, the copy numbers of *MET*, *HGF*, and *EGFR* progressed from a balanced conformation relative to ploidy state in the pre-treatment MPNST, to an imbalanced amplification above regional ploidy state in the recurrent MPNST (Figure 1D, Figure 2A). *MET* copy number gain was confirmed in the pretreatment and recurrent MPNST using quantitative copy number PCR (Supplemental Figure 1). Loss of *NF1* and *TP53* was also observed in the pre-, post-and recurrent-MPNST samples (Figure 1D, Figure 2B). Interestingly, the non-synonymous coding alteration in the *TP53* gene, NC_000017.11:g.7673223G>C, was found to be present in combination with *TP53* hemizygosity in all stages of the MPNST. A similar phenomenon was observed for a synonymous coding mutation at the hemizygous *CDKN2A* locus in the metastatic tumor. Copy neutral loss of heterozygosity events were observed in the post-treatment sample in the genes *CHD4*, *EED*, *EPC1*, *PTEN*, and *RASSF1*. These observations demonstrate an important role for *MET* and *HGF* in the earliest stages of MPNST disease progression.

**Figure 2:**
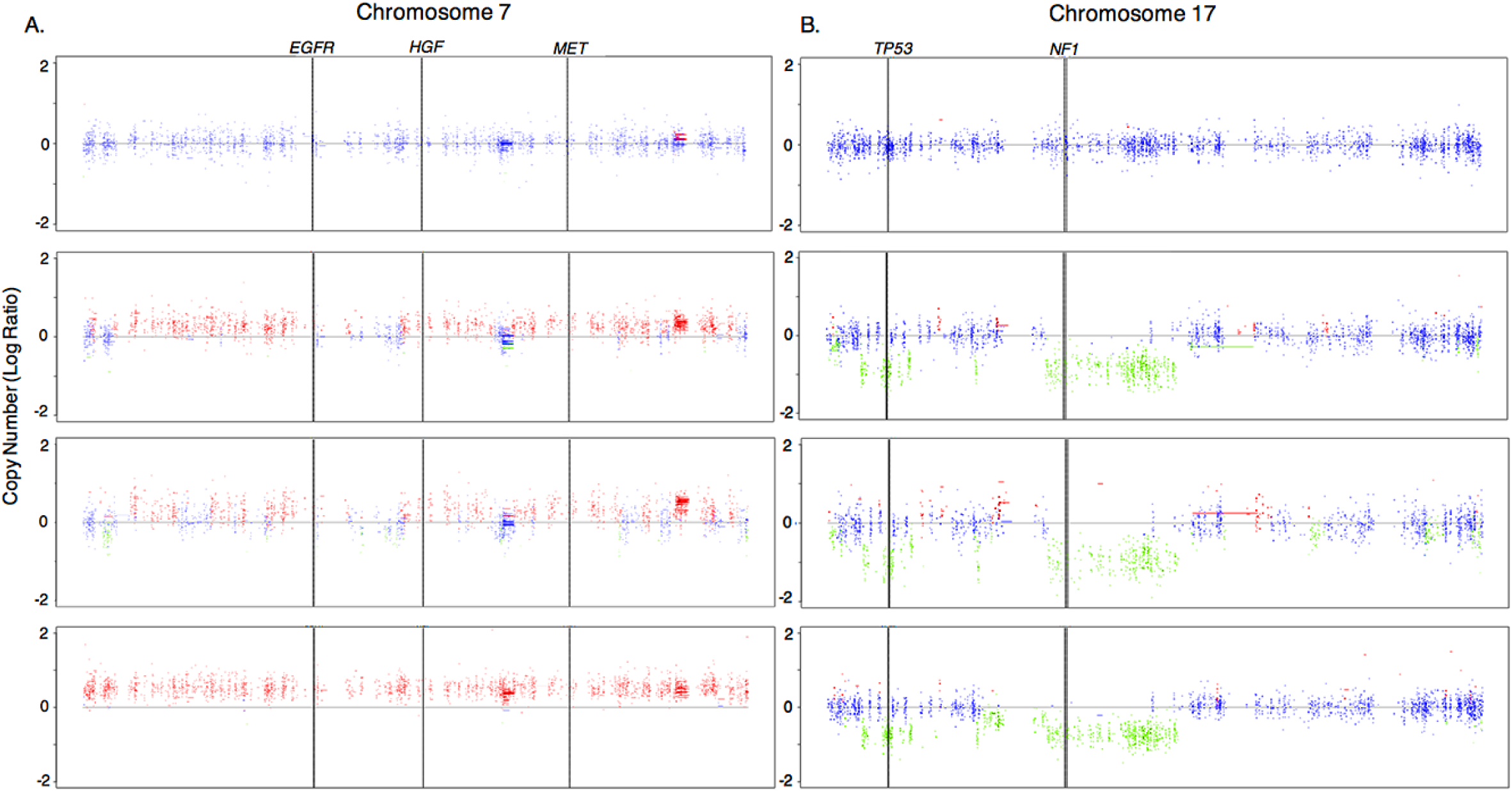
Gain of MET, HGF, EGFR and loss of NF1 and P53 copy number observed during MPNST progression. TITAN copy number analysis of variants found between blood and MPNST samples for each stage of progression was performed on A) Chromosome 7 and B) Chromosome 17. Blue = Neutral, Green = Hemizygous loss of heterozygosity, Red = Gain. (n=1 sample per time point).

### *Met* amplification and NF1 loss in peripheral nerve myelinating cells is sufficient to induce MPNST formation

To interrogate the role of MET activation in NF1-related MPNSTs, we developed a mouse model that reflects *MET* amplification in the context of *NF1* deficiency. In this Cre-inducible mouse model (Figure 3A), a humanized *MET* transgene that was previously shown to activate MET signaling above physiologic thresholds in neuronal tissue (33,34) was combined with *NF1* deficiency in myelinating Schwann cells (30,35). These two genetic events were induced immediately after birth using a tamoxifen-inducible *Plp-creERT* transgene (36) in mice with a global, constitutive loss of the *NF1* allele. Tamoxifen induction at postnatal days 1-5 was selected to best replicate plexiform neurofibromagenesis based on previous studies of *NF1*^*fl/KO*^;*Plp-creERT*^*tg/+*^ mice (30,31). Allele-specific PCR confirmed recombination of the loxp-flanked NF1 gene in peripheral nerves and the presence of the recombined Met transgene (Figure 3B). For brevity we refer to the *NF1*^*fl/KO*^;*lox-stop-loxMET*^*tg/+*^;*Plp-creERT*^*tg/+*^ mice as **NF1-MET** mice throughout the manuscript. NF1-MET mice and controls (*NF1*^*fl/KO*^;*Cre, NF1*^*fl/+*^;*Met;Cre,* and wild-type) were aged for up to 600 days to evaluate their tumor phenotypes. The tumor free survival of NF1-MET mice was significantly decreased relative to the other four mouse lines (all *FDR q* < 0.001, Figure 3C and Supplemental Table 1). The odds of NF1-MET mice being euthanized due to tumor burden was 14.58 times higher than *NF1*^*fl/KO*^;*Cre* mice (*FDR q=*0.0008, 95% CI [3.51, 60.59]) and 23.33 times higher than *NF1*^*fl/KO*^ (*FDR q=* 0.0008, 95% CI [4.1, 132.9]) (Supplemental Table 2).

**Figure 3:**
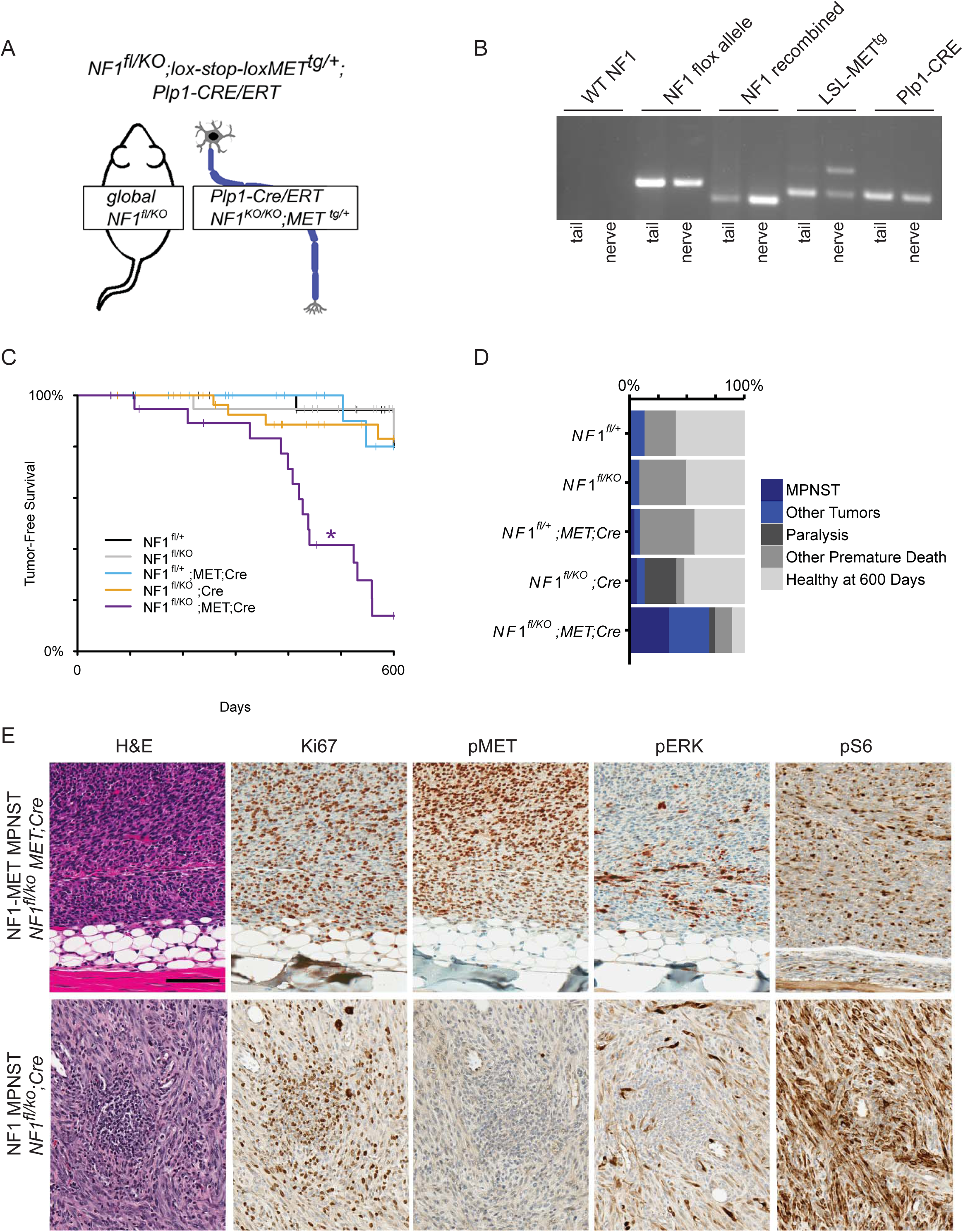
Met activation drives malignant tumorigenesis in a mouse model of NF1-related MPNST. (A) Schematic of mouse genetics. Mice have a global loss of one NF1 allele, and they have a loss of the second copy of NF1 plus activation of a MET overexpressing transgene in myelinating cells following tamoxifen-induction of Plp1-Cre/ERT activity at postnatal day 1-5. (B) Allele-specific PCR for a mouse with the combined NF1 loss and MET overexpression genotype shows absence of an unaltered NF1 allele (Wildtype NF1) and presence of the functional but loxp flanked NF1 allele (NF1 flox); the recombined, loss-of-function NF1KO allele (NF1 recombined); the inactive lox-stop-loxMET transgene (MET tg/rec, lower band) and the recombined tgMET transgene (MET tg/rec, upper band); and the Plp1-Cre/ERT transgene(Plp1-Cre). The first lane contains a DNA ladder. (C) Tumor-free interval data for all models plotted as Kaplan-Meier curves. NF1fl/KO;MET;Cre mice developed tumors significantly sooner than all other mouse lines (indicated by *). (D) Cause of death in the 600 day observation period is plotted by frequency. Blue indicates death or euthanasia related to tumor burden, greys indicate healthy after 600 days or death due to other causes. E) H&E staining confirmed MPNST histology (representative images shown of 6 samples assessed in triplicate). Ki67 verified high rates of proliferation. High pERK and pS6 were observed by immunohistochemistry, however high MET activation (pMET) was only present in the NF1-MET MPNST.

In total, 70% of NF1-MET mice developed neoplasms, 50% of which were MPNSTs; whereas in the 13% of control mice that developed neoplasms, none were MPNSTs (Figure 3C). MPNSTs derived from NF1-MET demonstrated the characteristic spindle cell morphology with a fascicular growth pattern (Figure 3D-E). Tumors were classified as MPNSTs according to the established Genetically Engineered Mouse (GEM) nerve-sheath tumor classification: Grade III, S100 +/ MyoD-with nuclear atypia, high mitotic rate, and focal necrosis or hemorrhage (Figure 3E)(37). Paralysis or pseudo-paralysis of the hind-limbs, occurring in 8 of 29 NF1-MET mice, was another major cause of premature euthanasia (Figure 3D, dark grey bars). Tumor initiation occurred earlier in NF1-MET mice than the other four mouse lines (all *FDR q* < 0.0001, Supplemental Table), with a median tumor-free survival time of 438 days (95% CI [408, 559]). Tumors from mice with activated MET transgene expression demonstrated a high rate of cellular proliferation (Ki67) and greater amounts of active phospho-MET (pMET) by immunohistochemistry (Figure 3E).

In order to evaluate MET signaling in other genomic contexts, we isolated MPNSTs from both *NF1*^*KO/+*^;*p53*^*LSL-R172H*^ and *NF1*^*fl/KO;*^*Plp-creERT*^*tg/+*^ mice. *NF1*^*KO/+*^;*p53*^*LSL-R172H*^ mice (referred to as **NF1-P53**) were derived by crossing the *NF1*^*KO/+*^ and *p53*^*LSL-R172H*^ mice (38). These mice did not require crossing with a *Cre* recombinase for tumor induction, since they are essentially a p53^KO/+^ model as the LSL cassette prevents expression of the *p53*^*R172H*^ mutant. MPNST tumors from NF1-P53 mice have LOH of the wildtype P53 allele and we confirmed that *p53*^*LSL-R172H*^ is in *cis* with *NF1* on Chr11. As a control for NF1 deficiency alone, we aged *NF1*^*fl/KO;*^*Plp-creERT*^*tg/+*^ mice and isolated an MPNST. For simplicity these mouse lines are referred to as **NF1** throughout the manuscript. Immunostaining of the NF1 MPNST showed less activated MET and ERK compared to the NF1-MET MPNST (Figure 3E).

### Copy number gains of MET and HGF observed in murine MPNSTs

Fluorescence *in situ* hybridization (FISH) was used to evaluate the *Rosa26 -* targeted *MET* transgene, the endogenous mouse *Met* gene, and the endogenous mouse *Hgf* gene. Confirmation of the transgene was performed with a R26-MET probe in NF1-MET mice. In normal spleen from a NF1-MET mouse, we observed 2-4 copies of the *MET* transgene and in MPNSTs we observed 2-8 *MET* transgene copies (Supplemental Figure 2 and Supplemental Table 3). We also performed FISH analysis to determine whether endogenous *Met* or *Hgf* were amplified in the MPNST tumors from the various NF1 mouse lines. We did not observe any additional endogenous mouse *Met* copy number gains in NF1-MET tumors (Figure 4A, Supplemental Table 4). Conversely, the NF1-P53 MPNST had a *Met* copy number gain of 2 in 50% of cells and the NF1 MPNST had a single *Hgf* copy number gain in 13% of MPNST cells (Figure 4B-C, Supplemental Table 4). To ensure the robustness of these models, each founder and the derived tumors were sequenced to verify both induced and spontaneous mutations.

**Figure 4:**
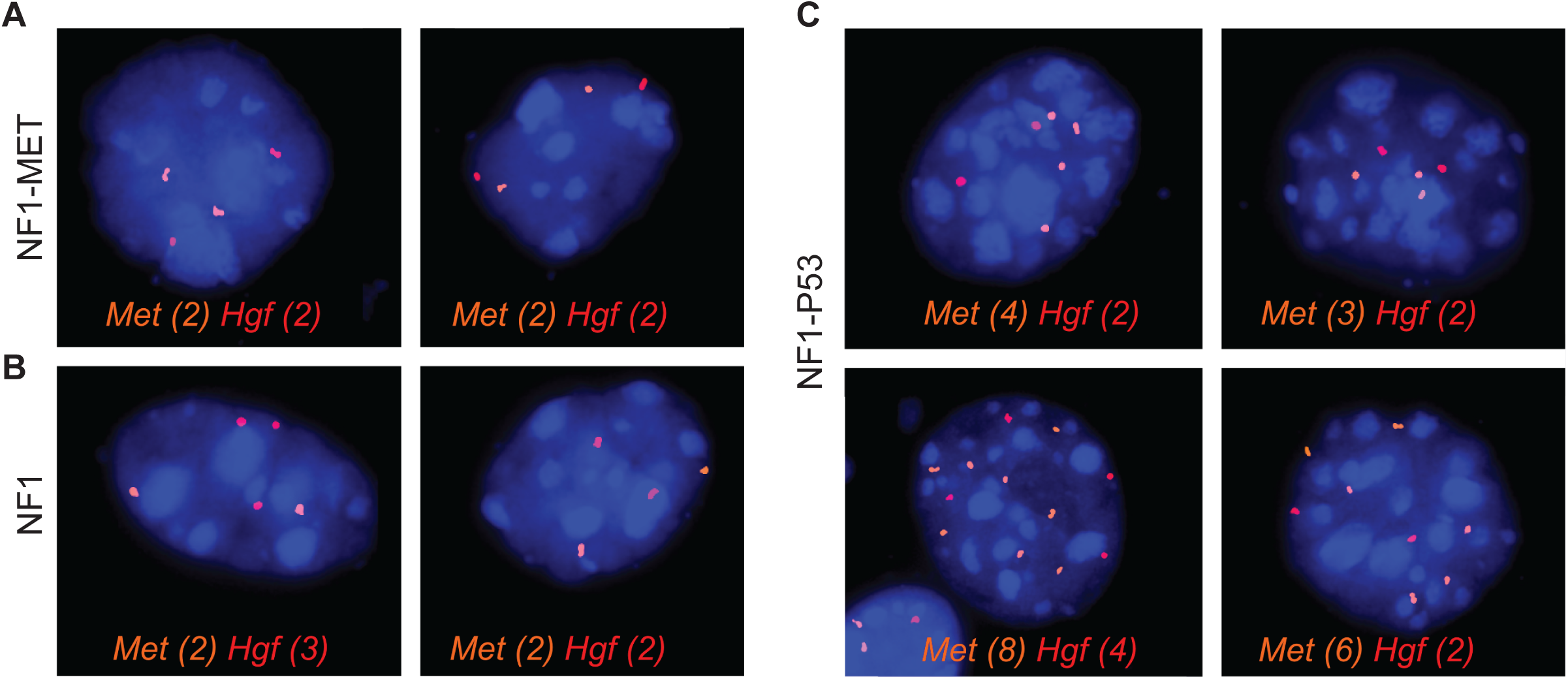
Interphase FISH demonstrates copy number gain of Met and Hgf in NF1-MET, NF1-P53, and NF1 MPNSTs. (A) No changes in endogenous Met (orange signal) and Hgf (red) copy number are observed in an MPNST from a NF1-MET mouse (MET transgene presence was confirmed, see Supplemental Figure 1). (B) Gain in endogenous Hgf copy number is observed in 12.5% of cells in a NF1 MPNST. (C) Gains in endogenous Met copy number is observed in an NF1-P53 MPNST with 4 copies in 49.5% of cells, 3 copies in 4% of cells, 6 copies in 5% of cells. An additional 6% of cells appear tetraploid with 8 copies of Met and 4 copies of Hgf. DAPI counterstain is shown in blue. Sample sizes and experimental details in Supplemental Table 3.

Additional genomic characterization of these NF1-MET mouse tumors, as well as tumors from control MPNSTs in NF1 and NF1-P53 mice was performed using 15-20x whole exome sequencing with the Agilent SureSelectXT2 mouse exome platform. Variants were cross-referenced against a list of genes with demonstrated roles in MPNST disease progression. No additional functional variants were detected in the NF1-MET, NF1, and NF1-P53 mouse models apart from the engineered genomic modifications. In summary, we observed endogenous Met or Hgf copy number gains in the NF1-P53 and NF1 tumors; however the NF1-MET MPNSTs had 4-10 copies of *Met* (transgene and endogenous Met) and represent a novel model of *Met -* amplified NF1-related MPNSTs.

### Differential kinome response to MET inhibition in murine MPNSTs

NF1 loss results in dysregulation of RAS-RAF-MEK signaling; however, studies in melanoma have demonstrated that NF1 has RAF-independent signaling activity (39). To investigate the effects of *NF1* loss, MET activation, and *P53* mutation in our MPNST models, we isolated stable cell lines from MPNSTs in NF1-MET, NF1-P53, and NF1 mouse lines. To examine the effects of MET inhibition on kinase signaling within each MPNST line, we used the kinase inhibitor capmatinib. Capmatinib (INC280) is an oral, highly selective and potent MET inhibitor that is well tolerated and has shown clinical activity in advanced solid tumors (40-42). Phase I and II clinical trials are ongoing to investigate the efficacy of capmatinib in cancers including melanoma, HCC, NSCLC, glioblastoma, colorectal cancer, and papillary renal cancer. To evaluate the effects of MET inhibition on downstream signaling in our distinct MPNST lines, we stimulated MET activation with HGF treatment and then treated with capmatinib (Figure 5). MET expression and activation was significantly higher in NF1-MET MPNST cells and required 5 nM INC280 to completely inhibit MET activation (Figure 5, left panel) whereas induced MET activation was eliminated with 1 nM INC280 in both the NF1-P53 and NF1 MPNST cells (Figure 5, middle and right panels). Only in the NF1-MET cells was ERK activation significantly inhibited by capmatinib, where in NF1-P53 and NF1 MPNST cells, ERK activation was marginally reduced by capmatinib treatment. Interestingly, HGF induced strong AKT phosphorylation in the NF1-P53 and NF1 cells (Figure 5, middle and right panels), yet in the NF1-MET cells AKT is highly expressed but minimally activated by HGF. When treated with 1-5 nM capmatinib, AKT activation is shut down in the NF1-MET and NF1 cells, but in the NF1-P53 cells pAKT is unaffected by MET inhibition. Each NF1 MPNST model has varying levels of MET activation that correspond with their sensitivity to MET inhibition. Our results suggest that NF1-depleted cells with MET amplification are highly dependent on MET and RAS/ERK signaling and AKT is likely not a signaling node of resistance. Interestingly, the increased AKT activation in capmatinib-treated NF1-P53 cells suggests this may be a unique mechanism of resistance to RTK inhibition in MPNSTs harboring *P53* loss or mutation. Therefore, MET inhibition may be an effective therapeutic strategy when a state of MET-addiction exists.

**Figure 5:**
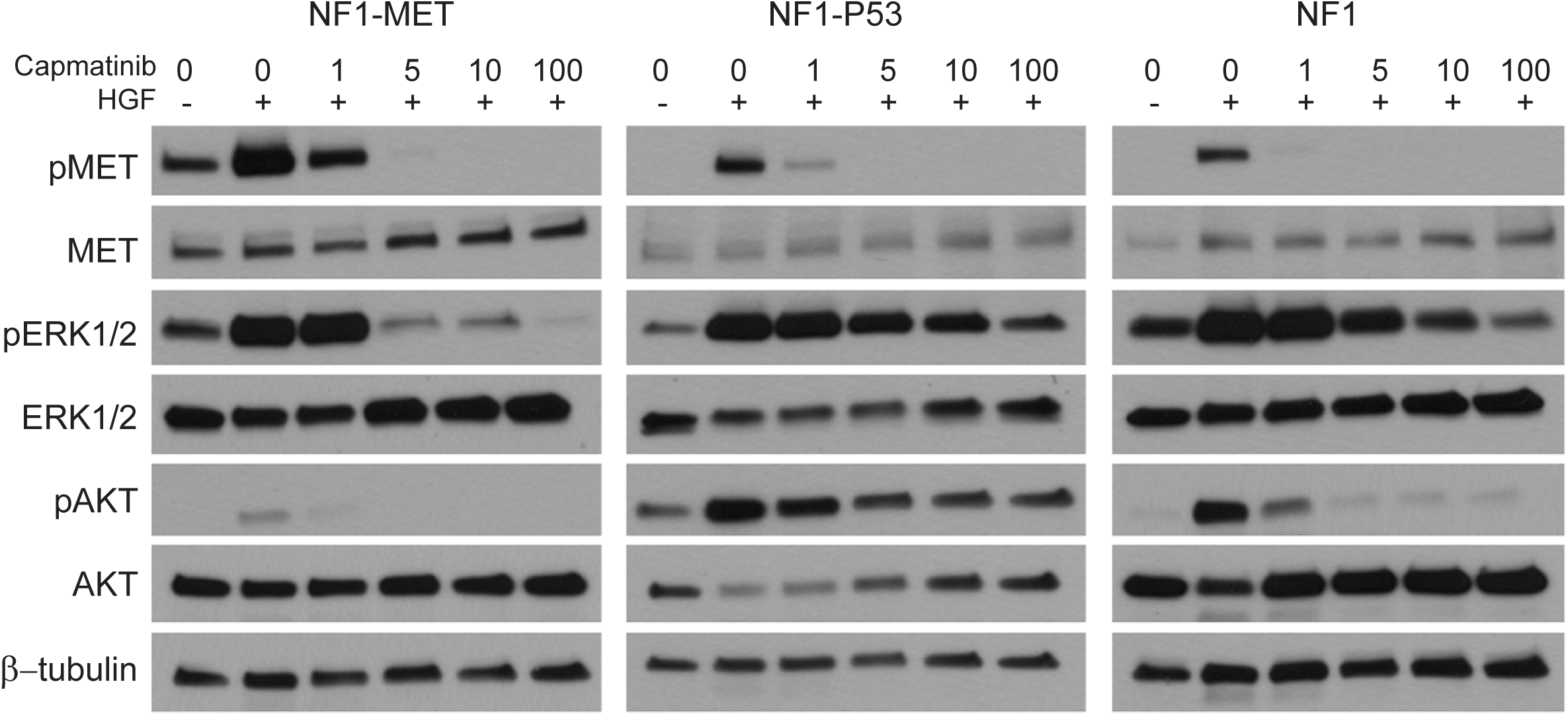
Response to MET inhibition of NF1-MET, NF1-P53, and NF1 MPNSTs. MET activation (pMET 1234/1235) and pERK1/2 (T202/Y204) areas inhibited by 5 nM INC280 in NF1-MET MPNSTs. ERK signaling was unaffected by INC280 treatment in NF1-P53 and NF1 tumors. Cells were serum starved overnight, then treated with INC280 for 2 hours. HGF (100ng/mL) was added for 15 minutes before harvesting lysates. Representative images shown for 3 samples assessed (westerns were repeated in triplicate).

### NF1-MET MPNST tumorgrafts are exquisitely sensitive to MET inhibition

Because MET is constitutively active in NF1-MET MPNSTs as compared to NF1-P53 and NF1 MPNSTs, we aimed to determine the effect of the MET inhibitor capmatinib on *in vivo* tumor growth. Tumorgrafts were established in immunocompromised mice from primary MPNSTs in the NF1-MET, NF1, and NF1-P53 models. Allele-specific PCR confirmed maintenance of the recombined NF1 gene and *Met* transgene. We tested the efficacy of three different concentrations of capmatinib (3, 10, and 30 mg/kg) on MPNST tumor growth. Capmatinib significantly inhibited tumor growth in the NF1-MET-derived tumorgraft model (Figure 6A-B, top panel). Once daily 3 mg/kg capmatinib significantly reduced the tumor growth rate by an average of 54.3 mm^3^/day (*FDR q <* 0.001, 95% CI [29.7, 78.9]) and 10 mg/kg dosage further reduced growth rates by an average of 56.1 mm^3^/day (*FDR q <* 0.001, 95% CI [34.0, 78.3]). Increasing the dose to 30 mg/kg did not produce a significant reduction, but did produce a more consistent effect (Figure 6A). By examining the individual growth curves for each tumor in the treatment compared to vehicle control groups, we assessed the variability in treatment response. The standard deviation of the 30 mg/kg dosage at 10 days was 181.5, compared to 228.4 for the 10 mg/kg treated mice and 465.6 for the vehicle treated mice (Figure 6A, top panel).

**Figure 6:**
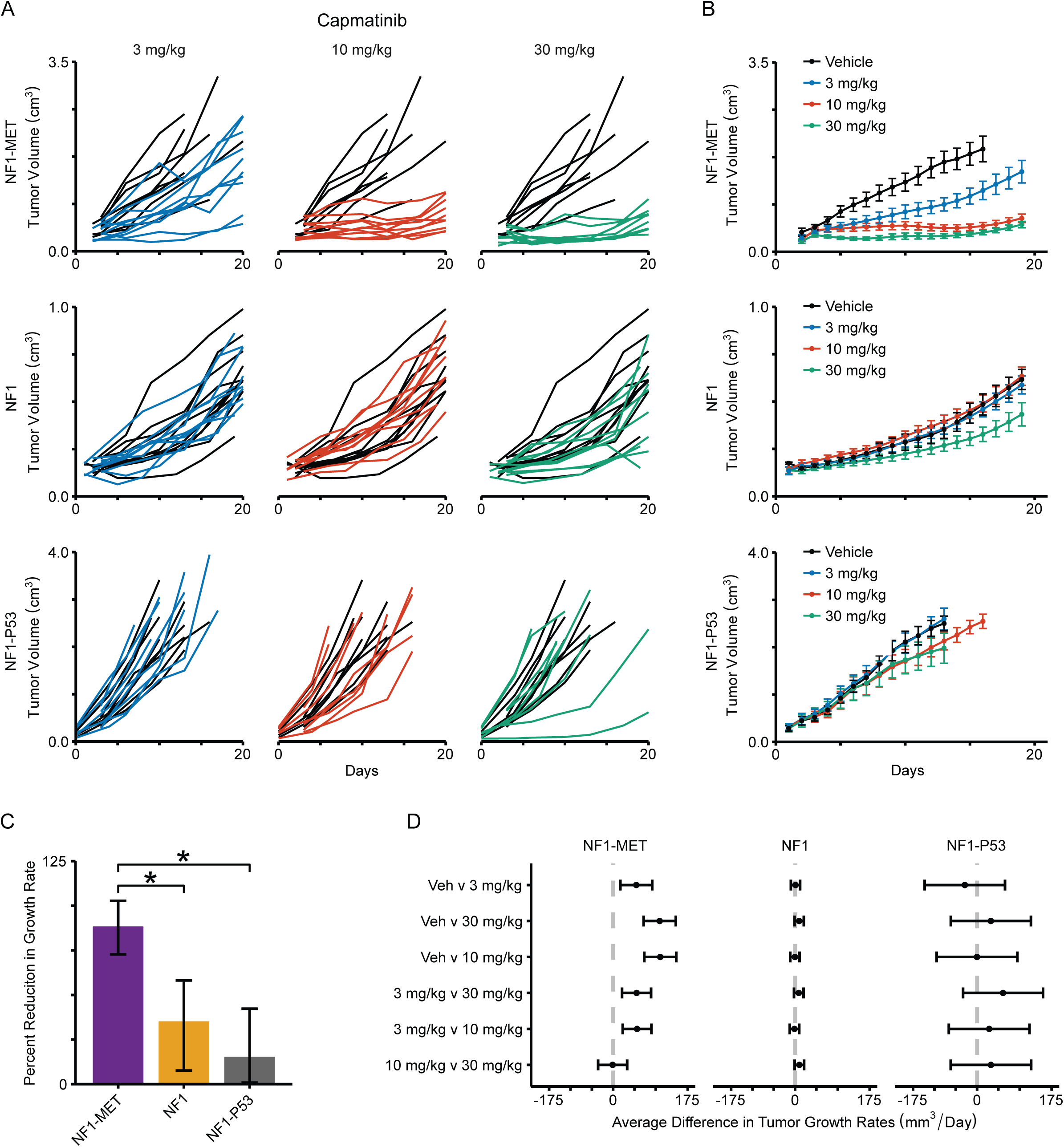
Tumorgrafts are differentially sensitive to MET inhibition with capmatinib. (A) Individual tumor growth plotted by line and treatment over 20 days. (B) Spline interpolated means and standard errors are plotted at each timepoint (N=8 mice per group). Curves terminate once >70% of mice have been euthanized in the respective treatment group. (C) Mean percent reduction in growth rates of mice treated with 30 mg/kg of capmatinib versus vehicle. Error bars are 95%, Bonferroni adjusted false coverage intervals, estimated via Fieller's theorem. (D) An * indicates non-overlapping confidence intervals between the three treatment groups. (E) Estimated mean differences between all treatments with 95% false coverage intervals by comparison and PDX line. Intervals are plotted on the same scale to highlight the differential tumor growth rates.

Capmatinib treatment of NF1 (Figure 6A-B, middle panel) and NF1-P53 (Figure 6A-B, bottom panel) MPNSTs did not show a significant response due to the heterogeneous treatment response; however in both the NF1-P53 and NF1 lines, we did observe a small subgroup of tumorgrafts that responded to 30 mg/kg capmatinib. The use of non-parametric bootstrap resampling revealed significant evidence that 30 mg/kg capmatinib reduced growth of NF1 tumors by an estimated 34.9% (Figure 6C, Bootstrap 95 percentile interval [7.62, 58.16]; Supplemental Table 4). Non-parametric bootstrapping also showed that capmatinib had the largest effect on NF1-MET MPNSTs with an almost 90% reduction in tumor growth rate relative to vehicle treatment (Figure 6C, Bootstrap 95 percentile interval CI [72.7, 102.7]; Supplemental Table 5). These results demonstrate the significant efficacy of MET inhibition in NF1-related MPNSTs with high MET expression and activity. Moreover, the results of capmatinib treatment in the NF1-P53 and NF1 lines suggest that MET inhibition is effective in reducing growth rate in a subset of tumors, yet combination therapy may be a more effective approach in MPNSTs with these genomic backgrounds.

## Discussion

The lack of effective therapies for MPNSTs remains a significant issue for individuals affected by NF1 (5-8). Our purpose in examining the evolution of genomic alterations in the case of a single, NF1-related human MPNST is twofold. Our first objective was to determine the degree of genomic instability that occurs over the course of MPNST treatment following malignant transformation. Intertumoral genomic heterogeneity is well documented in NF1-related MPNSTs (19,20,23,24,43), however expression profiling studies do not necessarily support molecular subtyping at the expression level (23). One theory is that an initial catastrophic event drives somatic and structural variation around the time of malignant transformation, but the genome remains relatively fixed over time. Evidence supporting this idea can be found in the characterization of human MPNST PDX models that demonstrate consistent genomic stability throughout expansion and engraftment (44). Our second objective was to identify targetable drivers of disease progression in MPNSTs. Somatic and structural variants that appear early and accumulate throughout disease progression represent de facto driver mechanisms and in the case of NF1-related MPNSTs, could indicate fundamentally important signaling pathways in Schwann cell dedifferentiation (45,46). Both findings may highlight susceptible pathways or biomarkers for drug targeting.

In our analysis, we identify in the pre-treatment specimen hemizygous microdeletions in the *NF1* and *TP53* loci, with concomitantly observed amplifications of *MET*, *HGF*, and *EGFR*. It is not surprising that a *TP53* variant was present in the pre-treatment MPNST given the prevalence of *P53* mutations in clinical samples (47), and the known driver effects of *P53* in the context of *NF1* deficiency (48); however, the early concomitant presence of *MET*, *HGF*, and *EGFR* amplifications, and the site-specific expansion of these loci over the course of treatment points to an adaptive mechanism for both malignant transformation and clonal selection. The relationship between *P53* haploinsufficiency and *EGFR* amplification has been previously demonstrated in human MPNSTs (49); however the timing and the relationship between *MET/HGF* and *P53* genomic status has not been evaluated. Direct cooperation between MET and RAS has been shown to promote tumor resistance in other cancers. For example, KRAS and MET amplification mediates acquired resistance to MET inhibitors in MET-addicted gastric and lung cancer cells (50). In an esophagogastric cancer patient treated with a MET inhibitor, a KRAS mutation was discovered as a novel cause of acquired resistance (51). Whereas in KRAS-addicted cancer cells, KRAS mediated MET expression via increased MET translation and promoted “KRas addiction” in anchorage-independent conditions (52). Given the pleiotropic effects of MET activation (25,26) and its emerging role in therapy resistance (53-56), these data provide strong rationale for further exploration of MET-RAS and RAS-MET signal interactions in NF1-related MPNSTs.

Even though therapeutically targeting RTKs has been successful in other cancers, the failure of EGFR inhibition (erlotinib) in MPNSTs indicates that other RTK pathways should be evaluated (57). We hypothesized that MET activation is an early and sufficient event for malignant transformation in NF1-related MPNSTs, and a bona-fide therapeutic target in a subset of “MET-addicted” MPNSTs. To interrogate the role of MET activation in NF1-related MPNSTs, we developed a novel mouse model that reflects MET activation in the context of NF1-deficient Schwann cells. In addition to plexiform neurofibroma formation, NF1-MET mice express a robust MPNST phenotype in the absence of additional spontaneous or induced mutations. An interesting observation is that MPSNT tumorgrafts derived from NF1-P53 mice exhibit spontaneous *Met* amplification and MPNST tumorgrafts derived from NF1 mice exhibit spontaneous *Hgf* amplification. FISH analysis revealed *Met* and *Hgf* copy number variation within tumors supporting the concept that locus-specific instability exists in murine MPNSTs. Despite the presence of *Met* and *Hgf* amplifications respectively, there was no evidence of constitutive MET activation in the NF1-P53 or NF1 models, whereas the NF1-MET tumors were strongly activated at baseline. MET activation was further induced with HGF treatment in all of MPNST tumorgraft cell lines. Collectively, these data confirm that the degree of MET activation is dependent on the genomic context and that MET activation is sufficient for malignant transformation in NF1-deficient Schwann cells to MPNSTs in the setting of germline NF1 haploinsufficiency.

To investigate the therapeutic potential of MET inhibition among our series of genomically diverse tumorgraft models, we chose the highly selective MET inhibitor, capmatinib which is currently being tested in the setting of EGFR therapy resistance in NSCLC. We demonstrate that MET-activated (NF1-MET) MPNSTs were uniformly sensitive to capmatinib therapy whereas Met-amplified NF1-P53 and Hgf-amplified NF1 tumors were only partially inhibited due to response heterogeneity within individual treatment groups. These data represent the first demonstration of capmatinib treatment response in NF1-related MPNSTs. It follows that when MET activation occurs a priori, the subsequently derived MPNSTs exhibit a MET-addicted signature whereby both RAS-ERK and PI3K-AKT activation are completely suppressed by selective MET inhibition. These findings further support the concept that MET inhibition is a viable treatment option for the subset of MPNSTs bearing a MET-addicted signature. Currently, it is difficult to define the size or makeup of the “MET-addicted” clinical subset in the NF1 population due to the lack of suitable biomarkers, however MET Y1234/35 phosphorylation has been proposed as a candidate immunohistochemical marker to define MET-activated MPNSTs, and predict MET inhibitor response (29).

As discussed previously, we observed a heterogeneous treatment response to capmatinib in the NF1-P53 and NF1 tumor groups. One explanation for this partial response is that MPNSTs maintain a fundamental plasticity for reprogramming of RAS effector kinase signaling in the setting of MET inhibition similar to what is observed in breast, lung, and gastric cancers (62-66). The majority of RTKs drive RAS/ERK and PI3K/AKT/mTOR signaling therefore resistance to MET inhibition may be mediated through alternative RTKs that are recruited to support RAS/ERK and PI3K/AKT signaling such as EGFR, PDGFR or both. Alternatively, the emergence of secondary genomic alterations could also drive selection of resistant cell populations. In the NF1-P53 murine MPNST model, EGFR overexpression occurs as a result of locus-specific amplification (58). EGFR amplification could partially mitigate response to capmatinib similar to what is observed in MET-addicted gastric carcinoma cells treated with MET inhibitors (59). We show that HGF-inducible MET activation is fully inhibited by capmatinib in all tumor models; however, persistence of downstream RAS/MAPK and PI3K/AKT activation occurred in both the NF1-P53 and NF1 models in the setting of complete MET inhibition. Both ERK and AKT activation persisted or was even amplified at the highest concentrations of capmatinib as a result of increased protein synthesis to sustain phosphorylation-dependent activation. These results suggest that the modulation of capmatinib response was dependent on the genomic context of the models. In our companion study Pridgeon et al., we examined how the genomic status of MET and P53 affects the efficacy and kinome response of kinase inhibitors in MPNSTs (60). Overall, these results highlight the challenges of treating genomically diverse MPNSTs with single agent tyrosine kinase inhibition strategies, as well as highlight the diverse mechanisms that promote MPNST progression and therapy resistance.

## Methods

### Whole genome exome sequencing and TITAN analysis

Extracted DNA samples for both human and mouse were prepared using Agilent SureSelect library prep with the Agilent SureSelect Human All Exon V5 and Agilent SureSelectXT2 exome capture systems, respectfully. Both human and mouse exome samples were sequenced with 2X 100bp reads on the Illumina HiSeq 2500 from the Michigan State University Research Technology Support Facility (MSU-RTSF) with a total mean read coverage of 50-90X and 15-20X, respectively. Read quality was assessed using FASTQC (http://www.bioinformatics.bbsrc.ac.uk/projects/fastqc/) and trimmed using seqtk (https://github.com/lh3/seqtk). After trimming, reads were mapped to the hg19 genome using BWAMEM v 0.7.12 (http://arxiv.org/abs/1303.3997) and duplicates were removed with Picard MarkDuplicates (http://broadinstitute.github.io/picard/). Variant calling using the deduplicated BAMs was completed using GATK Best Practices with suggested default values (61-63).Somatic variants were identified using MUTECT2 (64) using default settings. Copy number alterations (CNA) of MPNST samples were assessed using TITAN (65) for pre, post, and recurrent samples compared to blood exome. Default ploidy values were set to two and converge parameters were set to default values. Segment calls generated by TITAN were used for both genome wide and inferred gene specific copy number alternation analyses.

### Development of NF1-MET, NF1-P53, and NF1 mice

NF1^fl/KO^;lox-stop-loxMET^tg/+^;Plp-creERT^tg/+^ mice and controls were created to examine the combination of conditional NF1 loss of heterozygosity (42, 43) with overexpression of a humanized *MET* transgene (33,34). These two genetic events were targeted to occur in myelinating cells immediately after birth using a tamoxifen-inducible *Plp-creERT* transgene (44) in mice with a global, constitutive mutation of the other allele of *NF1*. *NF1*^*fl/+*^ mice were obtained from the NCI Frederick repository. *NF1*^*KO/+*^ mice were created by breeding *NF1*^*fl/+*^ mice to *CMV-Cre* mice (66). Only second generation or later *NF1*^*KO/+*^; *Cre*-negative animals were used for subsequent breeding. *Plp-creERT* (B6.Cg-Tg(Plp1-cre/ERT)3Pop/J) and *CMV-Cre*(BALB/c-Tg(CMVcre)1Cgn/J) transgenic mice were obtained from The Jackson Laboratory. *NF1*^*fl/KO*^;*lox-stop-loxMET*^*tg/+*^;*Plp-creERT*^*tg/+*^ mice were produced by crossing *NF1*^*fl/fl*^;*Plp-creERT*^*tg/+*^ mice with *NF1*^*KO/+*^;*lox-stop-loxMET*^*tg/+*^ mice. Lactating dams were dosed with 100 µl of 20mg/ml tamoxifen (Sigma) in corn oil (Acros Organics) by oral gavage b.i.d. on postnatal days 1-5, based on similar previous studies (31). Of note, crosses using *NF1*^*fl/fl*^;*Plp-creERT*^*tg/tg*^ mice did not yield pups at expected Mendelian ratios and were not used for this study. Nf1^*KO/+*^ mice were created by breeding *NF1*^*fl/+*^ mice to mice carrying a *CMV-Cre* transgene. Only second generation or later *NF1*^*KO/+*^; *Cre -* negative animals were used for subsequent breeding. *NF1*^*KO/+*^;*TP53*^*KO/+*^ mice were created by breeding *NF1*^*KO/+*^ mice to 129S4-*Trp53*^*tm2Tyj*^/Nci mice (functionally *TP53*^*KO/+*^) (38). All *NF1*^*KO/+*^;*TP53*^*KO/+*^ mice produced were bred to wildtype animals to confirm *cis*-conformation of *NF1* and *TP53* as performed in Vogel et al. (67). Mice producing at *NF1*^*KO/+*^;*TP53*^*KO/+*^ offspring when bred to wild-types were used as founders. All animal experimentation in this study was approved by the Van Andel Institute's Internal Animal Care and Use Committee. Details on mouse genotyping procedures are in Supplementary Data.

### Development of murine MPNST tum orgrafts and drug treatment

Immediately following euthanasia of tumor bearing mice, 15-25 mg portions of each tumor were transplanted into the flank of athymic nude mice using a 10 gauge trochar. Mice were examined weekly and euthanized when the tumor size exceeded 1500 mm^3^. For treatment studies, bulk tumor pieces were transplanted subcutaneously into athymic nude female mice and tumor growth was evaluated twice weekly. When tumor volume reached approximately 150 mm^3^, mice were randomized into vehicle, capmatinib 3 mg/kg, 10 mg/kg, or 30 mg/kg treatment groups and dosed 5 days per week via oral gavage for 3 weeks or until mice reached euthanasia criteria. Capmatinib was obtained from Novartis. The tumors were measured twice weekly using a caliper and the tumor volumes were calculated as length x width x depth.

### Statistical Methods

Kaplan-Meier curves were used to display tumor-free survival and, after verifying the proportional hazards assumption, Cox regression with false discovery rate adjusted contrasts were used to test for differences in tumor incidence rates between mouse lines. Three mice euthanized due to hindlimb paralysis had evidence of small paraspinal neoplasms at necropsy and were therefore included as tumor events. Genotypes with no tumor events are not shown. Logistic regression with false discovery rate adjusted contrasts were used to test for differences in the frequencies at which the different mouse lines were euthanized/died early due to tumor burden. Linear mixed-effects models with false discovery rate adjusted contrasts were used to estimate and compare tumor growth rates between different capmatinib dosages within each line. Fiellers's theorem with Bonferroni adjusted significance level was used to estimate the confidence intervals for the percent reduction in tumor growth rates for mice treated with capmatinib versus vehicle across the different lines (68). All analyses were conducted using R v3.2.2 (https://cran.r-project.org/) with an assumed level of significance of α = 0.05.

### Study Approval

Tissue samples were collected in accordance with established IRB approved protocols at both Spectrum Health and the Van Andel Institute. Written informed consent was obtained from study participants prior to inclusion in the study.

*Detailed Methods for genotyping, FISH analysis, histology, and western analysis can be found in the Supplemental Data.*

## Authors' Contributions

**Conception and design:** JD Peacock, CR Graveel, MR Steensma

**Development of methodology:** JD Peacock, EA Tovar

**Acquisition of data:** JD Peacock, MG Pridgeon, EA Tovar, J Koeman, J Grit, CJ Essenburg

**Analysis and interpretation of data:** M Bowman, Z Madaj, RD Dodd, DM Cardona, M Chen, DG Kirsch

**Writing, review, and/or revision of the manuscript:** JD Peacock, CR Graveel, MR Steensma

**Administrative, technical, or material support:** F Maina, R Dono

**Study supervision:** MR Steensma

## Acknowledgements

We would like to thank the MSU RTSF Genomics Core for genomic analysis of the MPNST models, Patrick Dischinger for technical assistance with this study, Amanda Schuiling for assistance with statistical analysis, and Anderson Peck from the VARI Small Animal Imaging Facility for his imaging support. We would like to thank Bryn Eagleson and the VARI Vivarium for their continuous dedication.

## Supplemental Tables and Figures Legends

**Supplemental Figure 1:**
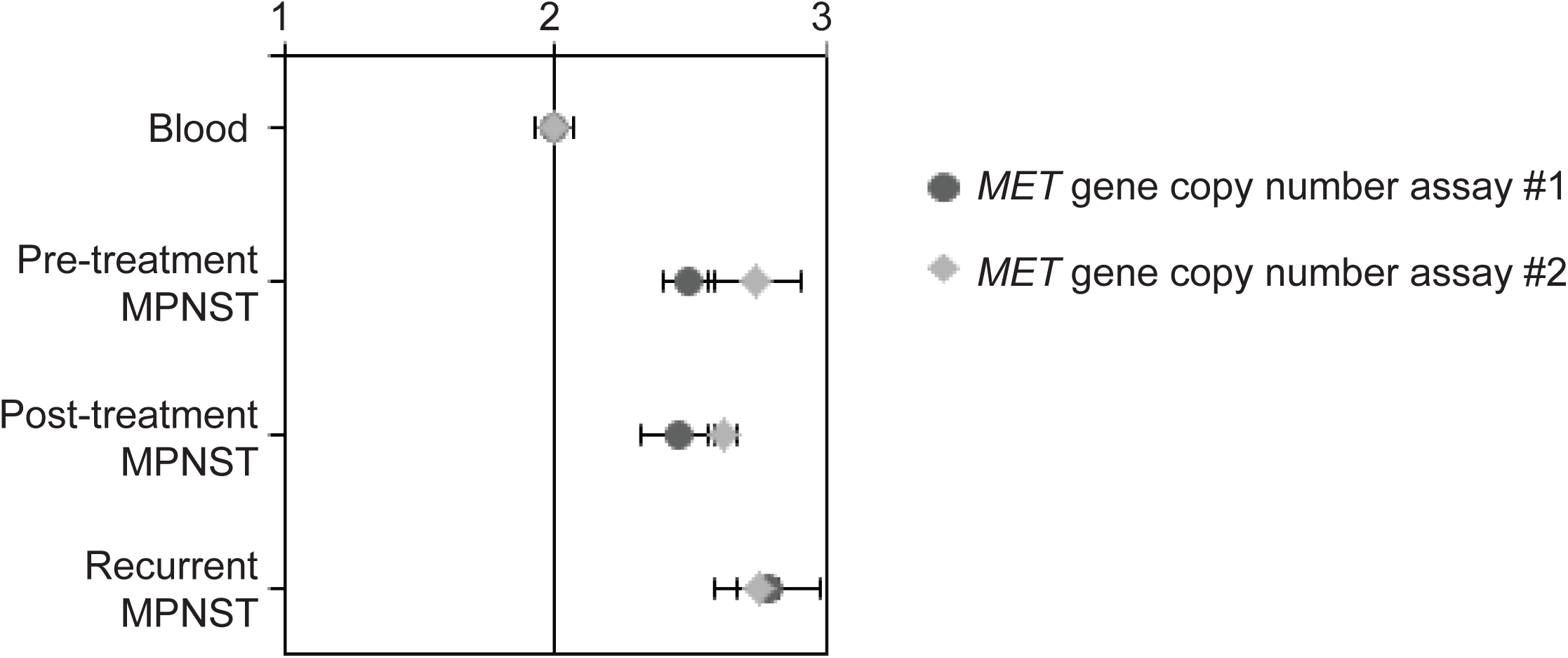
Increase in *MET* copy number during MPNST progression. Copy number changes in *MET* were confirmed with quantitative polymerase chain reaction using two assays. Average copy number is reported relative to blood DNA (N=2). Error bars indicate the standard error from technical triplicates.

**Supplemental Figure 2:**
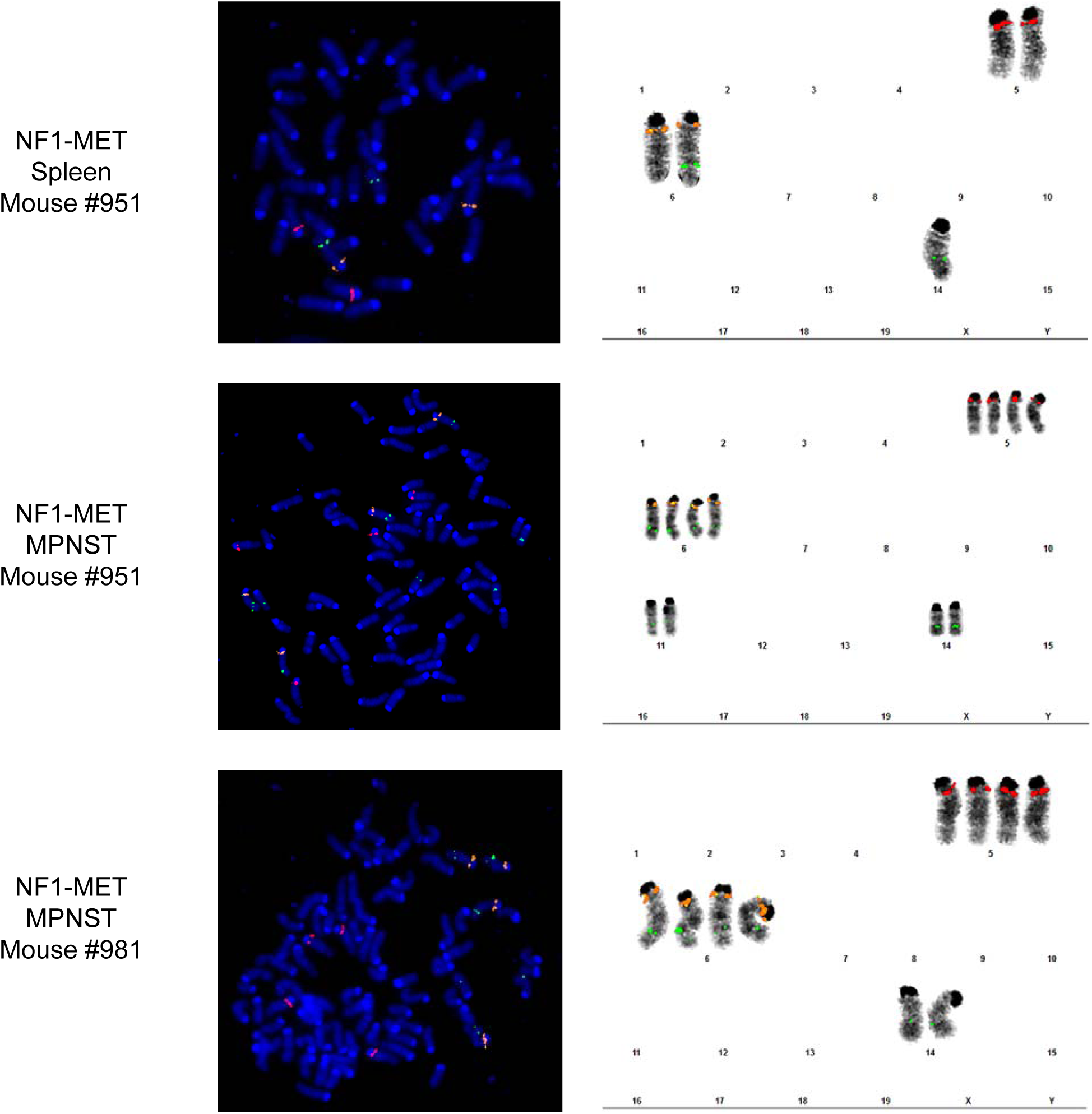
Gene copy number analysis of the *Met* transgene and endogenous *Met*. Metaphase FISH was used to measure the R26-MET transgene copy numbers. FISH probes r26-MET plasmid green, 6A2-1 (mouse Met) orange, and 5A2-2 (mouse HGF) red were used on normal spleen and MPNSTs from NF1-MET mice.

**Supplemental Table 1:** Cox Regression Analysis of Tumor-Free Survival. Results from testing for differences in tumor-free survival times between mouse lines. Mouse line *NF1*^*fl/KO*^;*MET;Cre* was used as the reference group. FDR q represents false discovery rate adjusted p-values.

| Line | Cox PH Coefficient | SE | P |
| --- | --- | --- | --- |
| *NF1<sup>fl/KO</sup>* | -2.3612 | 0.6489 | 0.0003 |
| *NF1<sup>fl/+</sup>;MET;Cre* | -2.6354 | 0.7619 | 0.0005 |
| *NF1<sup>fl/KO</sup>;Cre* | -2.3409 | 0.7614 | 0.0021 |
| *NF1<sup>fl/+</sup>* | -2.2420 | 0.5791 | 0.0001 |

**Supplemental Table 2:**
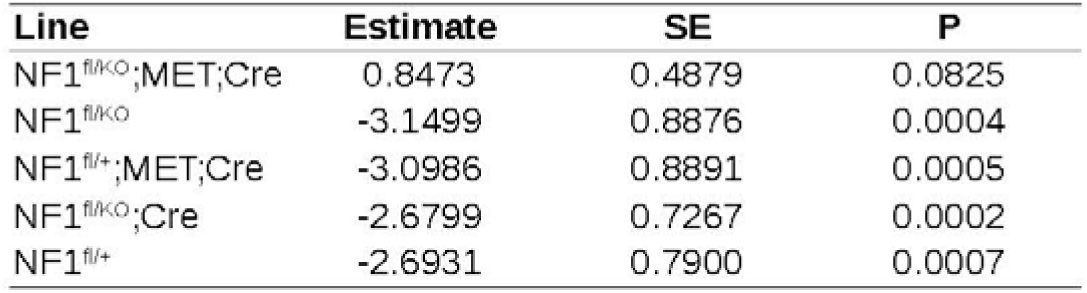
Logistic Regression Analysis of Cause of Death Frequency. Results from testing for differences in the proportion of mice that had deaths related to tumor burden. FDR q represents false discovery rate adjusted p-values.

**Supplemental Table 3:** Summary of R26-MET transgene localization. Interphase FISH analysis spleen and MPNST cells from NF1-MET mice was performed and the r26-MET (green) signal was assessed. Note that several tumors had diploid and tetraploid cells present.

| Sample | # of r26-MET plasmid FISH Signals | r26-MET plasmid Integration Sites (Approximation) | Chr Count and Ploidy Level |
| --- | --- | --- | --- |
| NF1-MET<br>spleen<br>Mouse #951 | 2 [2]<br>3 [16]<br>4 [2] | 6E, 14E1 [2]<br>6E, 6E, 14E1 [13] / 6E, 11B5, 14E1 [3]<br>6E, 6E, 11B5, 14E1 [2] | 40<2n> [20] |
| NF1-MET<br>MPNST<br>Mouse #951 | 3 [10]<br>4 [6]<br>6 [2]<br>8 [2] | 6E, 6E, 14E1 [10]<br>6E, 6E, 6E, 14E1 [2] / 6E, 6E, 11B5, 14E1 [4]<br>6E, 6E, 6E, 6E, 14E1, 14E1 [2]<br>6E, 6E, 6E, 6E, 11B5, 11B5, 14E1, 14E1 [2] | 39~40<2n>[16]<br>79~80<4n>[4] |
| NF1-MET<br>MPNST<br>Mouse #981 | 2 [1]<br>3 [10]<br>4 [3]<br>6 [6] | 6E, 14E1 [1]<br>6E, 6E, 14E1 [10]<br>6E, 6E, 11B5, 14E1 [3]<br>6E, 6E, 6E, 6E, 14E1, 14E1 [6] | 40~41<2n>[14]<br>80~82<4n>[6] |
| NF1-MET<br>MPNST<br>Mouse #981<br>Xenograft | 2 [3]<br>3 [16]<br>6 [1] | 6E, 14E1 [1] / 6E, 6E [2]<br>6E, 6E, 14E1 [16]<br>6E, 6E, 6E, 6E, 14E1, 14E1 [1] | 40~42<2n>[19]<br>82<4n>[1] |

**Supplemental Table 4:** Copy number analysis of *Met* and *Hgf* in murine MPNST models. Relative copy number of Met and HGF was measured using dual-color interphase FISH on tumor touch preps from NF1-MET, NF1-P53, and NF1 MPNSTs. A summary of the FISH signals for the 6A2-1 and 6A2-2 (mouse Met) orange, and 5A2-2 (mouse Hgf) red is shown.

| Chromosome | 6A2/5A2 | 6A2/5A2 | 6A2/5A2 | 6A2/5A2 | 6A2/5A2 | 6A2/5A2 | 6A2/5A2 | Total<br>Cells<br>Counted |
| --- | --- | --- | --- | --- | --- | --- | --- | --- |
| # of FISH signals | 2O/2R | 2O/3R | 3O/2R | 4O/2R | 6O/2R | 6O/4R | 8O/4R |  |
| Sample |  |  |  |  |  |  |  |  |
| NF1-MET<br>MPNST<br>Mouse #981 | 100.0% | - | - | - | - | - | - | 200 |
| NF1<br>MPNST<br>Mouse #986 | 87.5% | 12.5% | - | - | - | - | - | 200 |
| NF1-P53<br>MPNST<br>Mouse#7900 | 32.5% | - | 4.0% | 49.5% | 3.0% | 5.0% | 6.0% | 200 |

**Supplementary Table 5:** Linear Mixed-Effects Modeling of Tumorgraft Results. Results of linear mixed-effects models, by line, which were used to estimate and identify differences in tumor growth rates. *FDR q* is the false-discovery rate q value (i.e. adjusted p-value). *FCI* stands for false coverage interval and is the false-discovery rate adjusted analog to a confidence interval.

| Line | Comparison | Estimated<br>Difference in<br>Tumor<br>Growth Rates | Std Error | FDR q Value | 95% FCI<br>Lower | 95% FCI<br>Upper |
| --- | --- | --- | --- | --- | --- | --- |
| NF1-MET | Vehicle v 3 mg/kg | 54.289 | 14.191 | 0.000 | 78.863 | 29.715 |
|  | Vehicle v 10 mg/kg | 110.422 | 14.191 | 0.000 | 134.996 | 85.848 |
|  | Vehicle v 30 mg/kg | 109.252 | 14.309 | 0.000 | 134.030 | 84.474 |
|  | 3 mg/kg v 10 mg/kg | 56.133 | 12.781 | 0.000 | 78.265 | 34.000 |
|  | 3 mg/kg v 30 mg/kg | 54.963 | 12.912 | 0.000 | 77.322 | 32.604 |
|  | 10 mg/kg v 30 mg/kg | -1.169 | 12.912 | 0.928 | 21.189 | -23.528 |
| NF1 | Vehicle v 3 mg/kg | 0.968 | 4.124 | 0.901 | 11.848 | -9.913 |
|  | Vehicle v 10 mg/kg | -0.512 | 4.124 | 0.901 | 10.368 | -11.392 |
|  | Vehicle v 30 mg/kg | 9.559 | 4.125 | 0.061 | 20.442 | -1.324 |
|  | 3 mg/kg v 10 mg/kg | -1.480 | 4.222 | 0.901 | 9.659 | -12.619 |
|  | 3 mg/kg v 30 mg/kg | 8.591 | 4.223 | 0.084 | 19.733 | -2.551 |
|  | 10 mg/kg v 30 mg/kg | 10.071 | 4.223 | 0.061 | 21.212 | -1.070 |
| NF1-P53 | Vehicle v 3 mg/kg | -28.651 | 35.730 | 0.513 | 65.614 | -122.916 |
|  | Vehicle v 10 mg/kg | -0.187 | 35.857 | 0.996 | 94.413 | -94.787 |
|  | Vehicle v 30 mg/kg | 32.346 | 35.600 | 0.513 | 126.267 | -61.575 |
|  | 3 mg/kg v 10 mg/kg | 28.464 | 35.860 | 0.513 | 123.072 | -66.145 |
|  | 3 mg/kg v 30 mg/kg | 60.997 | 35.603 | 0.513 | 154.926 | -32.932 |
|  | 10 mg/kg v 30 mg/kg | 32.533 | 35.730 | 0.513 | 126.799 | -61.732 |

**Supplementary Table 6:**
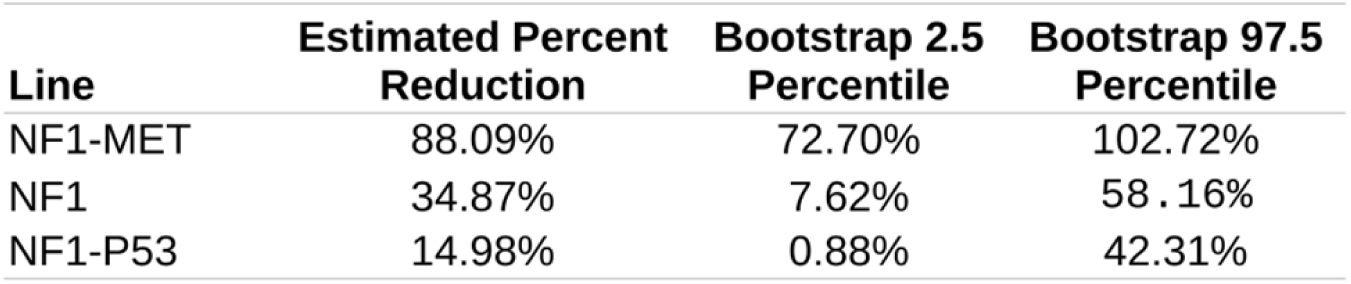
Bootstrap Resampling Results on Percent Reduction in Tumor Growth and Fieller Estimated Confidence Interval. Estimated percent reduction in growth rate, calculated as the mean growth rate of capmatinib 30 mg/kg normalized to the mean vehicle growth rate, by line. Fieller's theorem was used to calculate 95% confidence bounds, with Bonferroni adjustments for multiple testing.

